# Common multi-day rhythms in smartphone behavior

**DOI:** 10.1101/2022.08.25.505261

**Authors:** Enea Ceolini, Arko Ghosh

## Abstract

The idea that abnormal human activities follow multi-day rhythms spans ancient beliefs centered on the moon to modern clinical observations in epilepsy and mood disorders. Multi-day rhythms remain obscure in normal human activities. To address multi-day rhythms in healthy human behavior we analyzed over 300 million smartphone touchscreen interactions logging up to 2 years of day-to-day activities (N = 401 subjects). By using non-negative matrix factorization and data-driven clustering of ∼1 million periodograms, we captured a range of multi-day rhythms spanning periods from 7 to 52 days – cutting across age and gender. Despite their common occurrence, any given multi-day rhythm was observed in different parts of the smartphone behavior from one person to the next. There was little support in the data for ubiquitous rhythm drivers like the moon. We propose that multiple multi-day rhythms are a common trait, but their consequences may be uniquely experienced in day-to-day behavior.

## Introduction

Diurnal rhythms driven by intrinsic circadian clocks and environmental *zeitgebers* vividly influence our cognition and behaviour^1–4^. However, the role of multi-day rhythms is controversial across scientific disciplines and in society at large. In sleep research, evidence exists both supporting and refuting the idea that fluctuations in sleep duration follow lunar cycles^5–7^. In conventional human behavioral signals spanning reading ability to traffic accidents, multi-day rhythms are unconvincing^8–12^. Common notions on how the multi-day menstrual cycle impacts behavior is a source of workplace discrimination and extend to fundamental experimental designs ^13,14^. However, there is no evidence supporting that menstrual cycles make female behavior more susceptible to the 20 – 30 day rhythms than in males^15,16^. While the presence of multi-day cognitive and behavioral rhythms is unclear in the healthy population, it is clearer in certain clinical populations. In epilepsy, long-term data from brain implants and seizure diaries have helped reveal multi-day rhythms with 7, 15, 20 – 30-day periods^17–19^. In bipolar disorder, multi-day cycles are visible in mood fluctuations, which oscillate with 20 – 44, 54 – 59, and 80 – 89-day periods^20^. On the one hand, these longitudinal clinical observations raise questions about the origin of multi-day rhythms^21,22^. On the other, it remains unclear if multiday rhythms shape normal behavioral outputs in the real world.

The extensive log of human behaviors captured on personal computers offers a fresh avenue in the study of multi-day rhythms. Weekly cycles have been observed in the speed of smartphone interactions and the mood reflected in Twitter posts^23,24^. By leveraging the longitudinal smartphone interactions and the diverse behaviors engaged on the smartphone here we will address: (*i*) Which multi-day rhythms are present in human behavior in health? (*ii*) Are the rhythms determined by gender or age? (*iii*) Do the distinct rhythms influence different aspects of the behavior? By using behavioral periodograms we shall address multi-day rhythms well beyond what can be addressed by using the commonly used calendar structures such as the time of the day or the day of the week^25^.

To capture the behavioral diversity, we clustered the time series of smartphone touchscreen interactions (tappigraphy) according to the next-interval dynamics by using a joint interval distribution (JID)^26^. Here, the duration of an interaction interval *K* is considered in conjunction with that of the next interval *K* + 1 (**Fig. 1**). The probability densities of the joint intervals spanning 30 ms to ∼2 min is captured in 2500 two-dimensional bins. Periodograms derived from continuous wavelet transform of the time series of hourly JID can simultaneously probe a range of periods and behaviors (captured in the JID)^27^. The thousands of behavioral periodograms derived this way – from each individual – are difficult to interpret. Non-negative matrix factorization provides an interpretable form of dimensionality reduction where the analysis can be reduced to a handful number of *meta-rhythms* that are co-expressed across behaviors. The behaviors under the influence of the meta-rhythms can be identified using the reduced *meta-behavior* corresponding to each meta-rhythm. These meta-rhythms and behaviors are akin to genetic co-expression patterns, i.e., meta-genes in dimensionality reduction applied to genetic data or akin to facial features such as eyebrows in the reduction applied to facial images^28,29^. In this study, by using this approach we revealed the nature of multi-day rhythms spread across the diverse smartphone behaviors in a healthy population.

**Figure 1:**
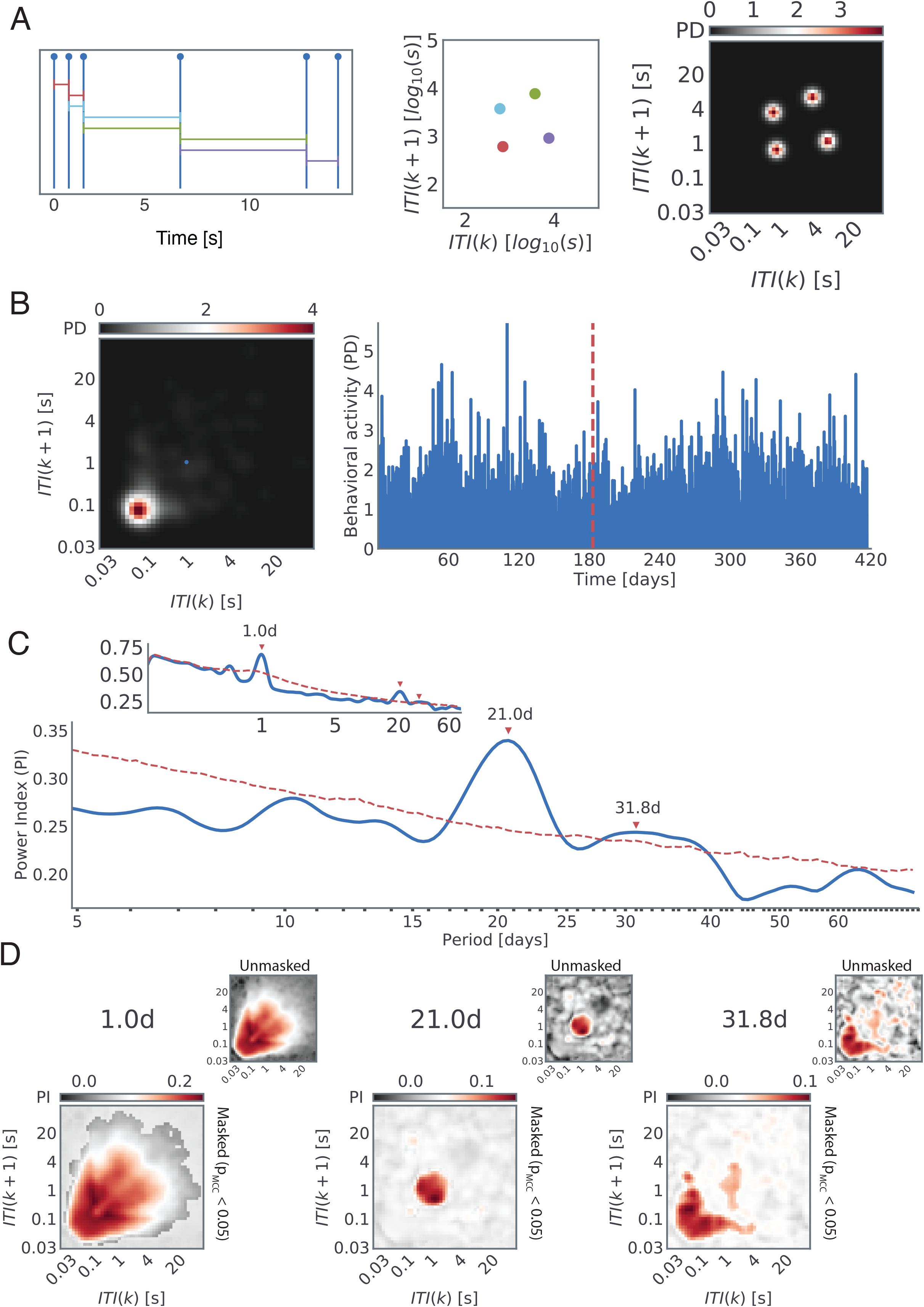
Spectral analysis of smartphone behavior based on inter-touch intervals. (**A**) We quantified smartphone behavior using the probability density of joint interval distribution (JID) in two-dimensional bins. An example of the probability density (PD) resulting from a series of 6 simulated interactions. (**B**) Example of behavioral activity of a subject captured by JID (left) accumulated over an hour-long window and (right) evolution of the probability density values at a select 2-dimensional bin over consecutive hourly windows (highlighted by using a small blue dot overlayed on the JID). (**C**) Periodogram for the two-dimensional bin selected above, obtained by averaging the continuous wavelet transform spectrogram over time, some of the peaks are marked using red arrows (red dashed line shows the 97.5^th^ percentile values based on block-bootstrap of the same data). (**D**) The power index (PI) of the selected periodogram peaks (red arrows in ‘C’) across the smartphone behavior. The two-dimensional bins that are not part of statistically significant clusters (multiple comparison correction, α = 0.05, ∼1000 block bootstraps) are masked with a translucent layer. The unmasked PI values are shown in the smaller inserts.

## Results

At the individual level, significant multiday rhythms could be established by using wavelet derived periodogram, in combination with the block-bootstrap method and 2-dimensional statistical clustering based on parametric *t*-tests ^27,30^ (**Fig. 1**). Examining the data at this level we observed that each multi-day rhythm appeared in a distinct set of smartphone intervals captured in the JID. Faced with such rich diversity we deployed non-negative matrix factorization to extract the co-expressed rhythms (*meta-rhythms*) and locate the behavioral repertoires in which the meta-rhythms were expressed (based on the *meta-behaviors* output of the factorization) (**Fig. 2**).

**Figure 2:**
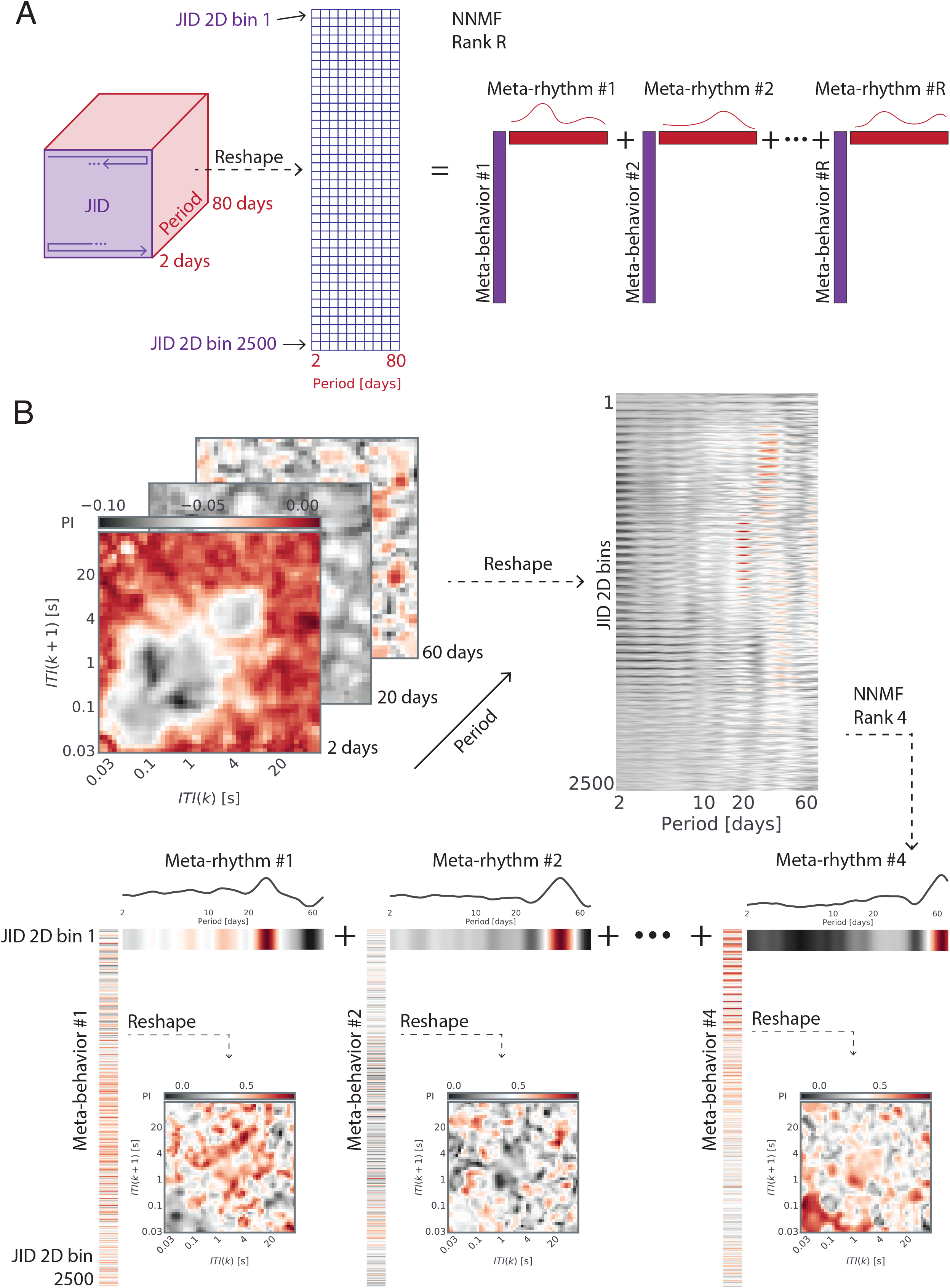
Non-negative matrix factorization reveals meta-rhythms and meta-behaviors. (**A**) A sketch of the three-dimensional tensor of periodograms across the behavioral space, and how it was reshaped to obtain a matrix of two-dimensional bins and periods. This matrix was decomposed by using non-negative matrix factorization and was operated on this matrix using optimal rank *R* (based on cross-validation) to obtain *R* meta-rhythms and the corresponding meta-behaviors. (**B**) An example of non-negative matrix factorization of rank *R* = 4 operated on the three-dimensional tensor of aperiodic component adjusted periodograms for one individual. In this subject, the meta-rhythms showed peaks at 27, 42, and 64 days, with the last peak being incomplete and such meta-rhythms with incomplete peak at the edge of the periodogram were not interpreted here. The meta-behaviors were reshaped into their original two-dimensional form to capture the distinct meta-behavioral expression corresponding to each meta-rhythm.

### Multi-day rhythms are invariant to age and cut across gender

The *meta-rhythms* were accumulated from across the population and clustered to yield 9 different meta-rhythms with peak periods of 7, 14, 19, 25, 32, 41 and 52 days (**Fig. 3**, for the 95^th^ percentile ranges see the values in ‘[]’ in the figure). The periods < 21 days were derived from recording lengths spanning a minimum of 90 days (N = 401, Supplementary Figure 1), whereas the periods > 21 days were derived from recordings spanning a minimum of 180 days (N = 218, Supplementary Figure 2). Aggregating across the population, the extracted meta-rhythms were largely dominated by a circa period rather than showing multiple prominent peaks. Essentially, the different multi-day rhythms are not simply co-expressed in the same sets of behaviors, as if they did then the factorization would have yielded meta-rhythms with multiple peaks. We established the prevalence of the meta-rhythms by counting their instances per person in the sampled population. According to the counts, the multi-day rhythms were present in the majority of the population – except for the 7-day rhythm – across the adult lifespan (**Fig. 3**). The 25-day meta-rhythm (95^th^ percentile peak width spanning 23 to 27 days) was exceptionally dominant in females compared to males – although 33% of the males did express this meta-rhythm.

**Figure 3:**
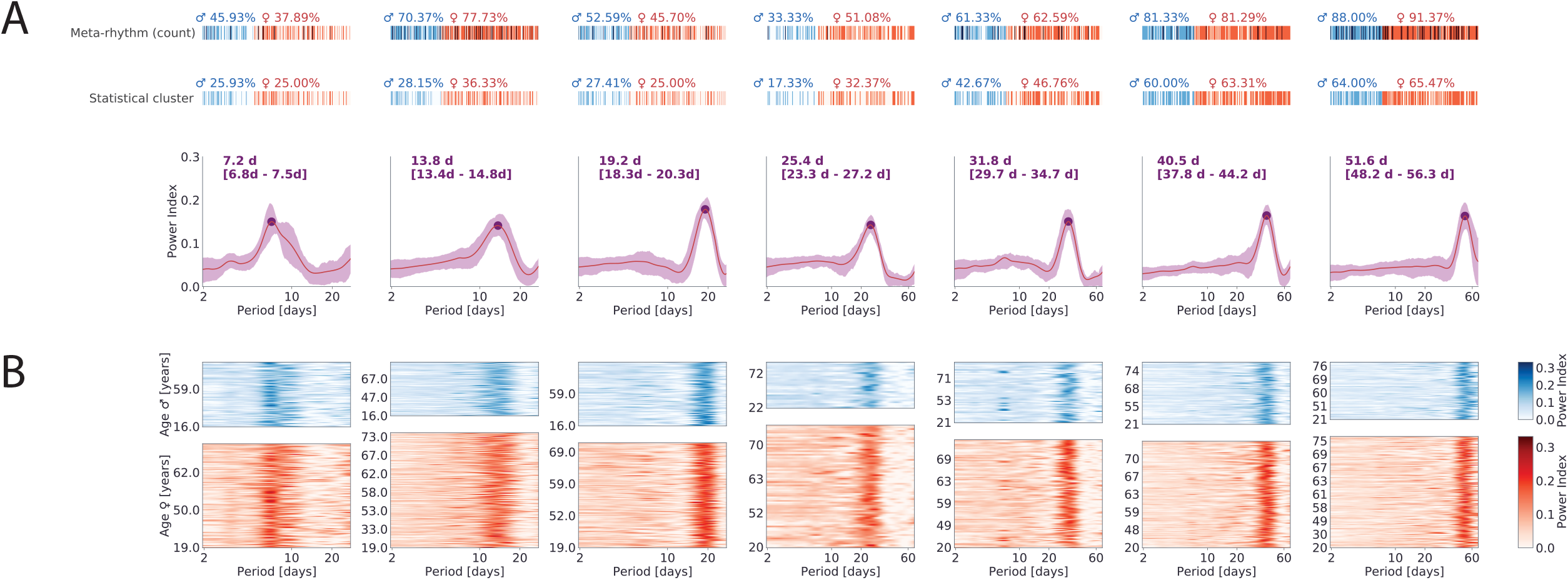
Clusters of common meta-rhythms in the sampled population, obtained with individualized optimal-rank non-negative matrix factorization. (**A**) Mean and the corresponding confidence interval (inter-quartile range, iqr, shaded) of meta-rhythms obtained from each cluster. The dot indicates the location of the average meta-rhythm peak (peak period noted above the plots along with the 95^th^ percentile range in ‘[]’). Each individual had zero or multiple meta-rhythms belonging to each cluster. This is indicated by the meta-rhythm count (first row) while the percentage indicates how many individuals (separately reported for males and females) present at least one significant statistical cluster (multiple comparison correction, α = 0.05, ∼1000 block bootstraps) at the identified multi-day rhythm. (**B**) Meta-rhythms assigned to each of the clusters identified in ‘a’, separated by gender and sorted by age.

In a follow-up analysis, we addressed the number of participants in whom the meta-rhythm-based periodogram peaks (95^th^ percentile width) were additionally part of statistically significant behavioral clusters based on block bootstraps of the JID time-series (**Fig. 3**) ^27,30^. Note, that this statistical clustering ignored rhythms that may be spread across highly diverse or non-neighboring behaviors on the JID. According to this approach < 30-day rhythms were present in minority of the population whereas the rest were present in either ∼50% of the population or higher.

### Multi-day rhythms are co-expressed across smartphone behavioral repertoires

Along with the meta-rhythms the factorization extracted the corresponding behavioral repertoires where the rhythms were expressed i.e., the meta-behaviors. To study these repertoires, we averaged the meta-behaviors across the people who showed significant rhythms according to the follow-up analysis described above (for average meta-behaviors without this selection see Supplementary Figure 3 and Supplementary Figure 4). The 7, 14, 19, and 32-day rhythms were preferentially expressed in the behaviors consisting of short consecutive intervals (**Fig. 4**). The 40 and 52-day rhythms were prominent across both the fast and slow behaviors. Interestingly, the 25-day rhythms were most prominent in the slow behavioral dynamics. These patterns were also present in the probability of the population showing a multi-day rhythm according to the parametric statistics (Supplementary Figure 3 and Supplementary Figure 4).

**Figure 4:**
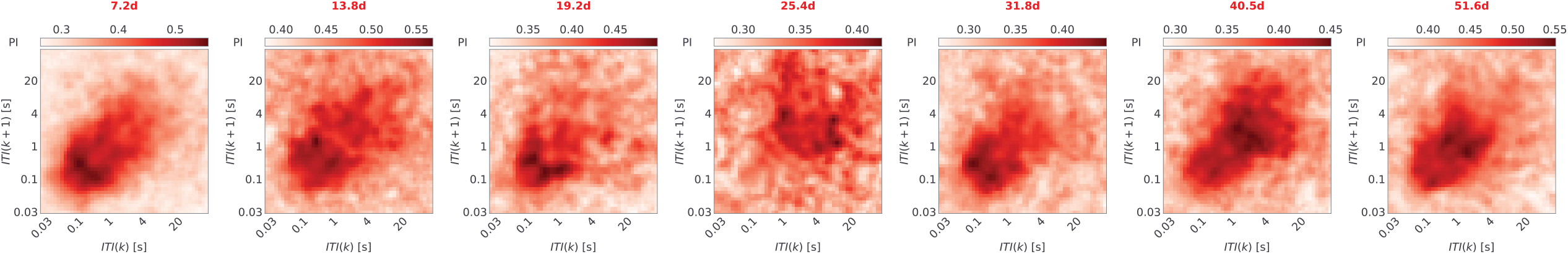
The expression of multi-day rhythms in the diverse temporal intervals captured on the smartphone. The non-negative matrix factorization and population level clustering revealed common multi-day meta-rhythms, and each meta-rhythm was accompanied by a corresponding meta-behavior. The mean meta-behaviors are shown for each of the identified multi-day rhythms. The mean was based on individuals where the identified rhythm was part of statistically significant clusters according to the parametric statistics based on block bootstraps.

### Sparsely coherent multi-day rhythms

A subset of the individuals (N = 175) was recorded from the same calendar window. We leveraged this to address whether the multi-day rhythms discovered here were synchronized according to common *zeitgebers* in this sample (**Fig. 5**). First, for a given subject and a given multi-day rhythm, we identified the two-dimensional JID bin with the maximum periodogram peak (Power Index) belonging to the largest statistically significant cluster. Next, we estimated the pair-wise phase coherence between individuals based on the identified bins. Unsurprisingly, when this approach was used on diurnal rhythms 99 % of the pairs were found to be phase coherent (Supplementary Figure 5). However, only a small fraction (< 30 %) of the pairs were phase-coherent for the multi-day rhythms with some of the rhythms being more likely to show coherence than the others. Thirty percent of the pairs at 7-day were coherent whereas < 10 % of the pairs were phase-coherent for the 14 & 19-day rhythms. Male-male, female-male, and female-female pairs did not substantially differ for any of the multi-day rhythms.

**Figure 5:**
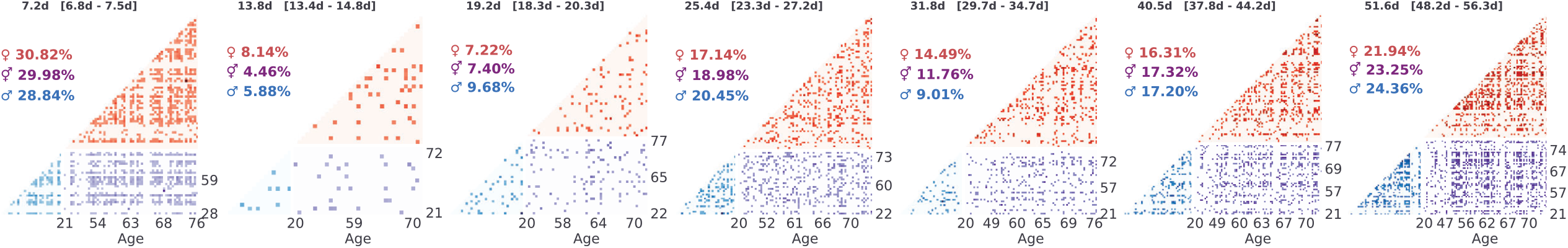
Limited spectral coherence across the population for the discovered multi-day rhythms. For each discovered multi-day meta-rhythm, the coherence across a population is shown in red for female-female, in blue for male-male, and in purple for male-female. Each lower-triangular matrix is sorted by age. From each individual, the JID 2-dimensional bin with the maximum power index from the largest significant cluster (according to parametric statistics based on block bootstraps) was chosen for pair-wise coherence analysis. The percentages indicate for each sub-group the rate of person-to-person statistically significant coherence (α = 0.05, ∼1000 bootstraps).

## Discussion

We extracted multi-day behavioral rhythms by using non-negative matrix factorization of thousands of periodograms that simultaneously probed a range of periods across the diverse behaviors on the smartphone at the level of each individual. While the weekly rhythms have been established in prior behavioral observations, they were less prevalent than the other rhythms discovered here spanning 14 to 52 days. The multi-day rhythms were present in a majority of the sampled population and were similarly prevalent across the adult life span and genders – with the notable exception of the 23 to 27-day rhythms (25-day meta rhythm peak, and henceforth referred to as ∼25-day rhythm) which was more prevalent in females.

All of the multi-day rhythms of ∼32 days or shorter that we report here have been previously reported in the studies tracking abnormal neural discharges captured by using brain implants in people with epilepsy^17,19^. In epilepsy – as in our behavioral observations in health – both males and females show ∼32-day rhythms ^31^. The ∼19, ∼25, ∼32, ∼41, ∼52-day rhythms have been observed in the mood fluctuations of patients with bipolar disorder^20^. Our findings raise the possibility that the multi-day rhythms in disease are driven by mechanisms already present in health ^22,22,31^.

One possibility is that the oscillators underlying the multi-day rhythms established here are driven by external *zeitgebers* such as the lunar cycles for the ∼32-day rhythms^21,32^. We reasoned that if the oscillators were simply driven by an environmental influence (such as lunar light), then the rhythms would be synchronized across the sampled population given the geographical constraint of the study (to The Netherlands). However, by using pair-wise phase coherence analysis we found no evidence for such mass synchronization for the multi-day rhythms. This does not rule out the role of lunar cycles in ∼32-day multi-day rhythm^21^. The Netherlands is highly urbanized with abundant artificial lighting, so the lunar light fluctuations could only have a negligible influence on the intrinsic clocks^5^. When out of reach of the lunar clues the intrinsic multi-day clocks may have turned free running. Alternatively, the oscillations may be driven by some unknown environmental factor or driven intrinsically such as in circadian genetic clocks ^2,33^. Less intuitively they may not be deterministic at all but emerge as an outcome of the complex interacting systems underlying behavior and the reception of environmental inputs even if the signal sources themselves are devoid of oscillations^34–38^.

The menstrual cycles do not easily explain the biased expression of the ∼25-day behavioral rhythm in females. After all, many of the males expressed the same rhythm, and the meta rhythm was found across the adult lifespan. In general, the resilience of multi-day rhythms even in advanced age indicates the role of fundamental underlying processes that do not succumb to the broad age-related behavioral, cognitive, and lifestyle changes. The behavior in which the multi-day rhythms were expressed varied from one rhythm to the next. For instance, the 25-day rhythm was expressed in the slow inter-touch interval dynamics while the 19-day rhythm was expressed in the fast dynamics. These differences indicate that there may be diverse mechanisms underlying multi-day rhythms resulting in distinct behavioral expressions.

Non-negative matrix factorization enabled us to interpret an otherwise complex expression of multi-day rhythms spanning diverse behaviors in healthy individuals and provides strong evidence for rhythmic real behavioral outputs that have long escaped scientific explorations. The behaviors in which these rhythms were expressed varied from person to person, and from one rhythm to the next. These variations may partly explain why multi-day rhythms have remained difficult to establish using conventional tools that only probe a select few behavioral features at a time. The presence of multi-day rhythms in real-world behavioral outputs propels the field forward to discovering their origins in health and establishing their role in diseases.

## Methods

### Recruitment and participants

Participants were recruited via on-campus advertisements, email, and the online data collection platform agestudy.nl. A Dutch subject registry (hersenonderzoek.nl) was used to approach participants to join agestudy.nl ^39^. This study pooled data across completed^23,26,40^ and ongoing data collections involving smartphone behavioral recordings in self-reported healthy participants (data collection frozen on the 2^nd^ of June 2022). The inclusion criteria were: (a) users with personal (unshared) smartphones with the Android operating system, (b) self-reported healthy individuals, (c) with no neurological or mental health diagnosis at the time of the recruitment, and (d) a minimum of 90 consecutive days of smartphone data. This resulted in 412 participants (age: min. 16 years, max. 84 years, median 59 years), and 403 reported their gender (64% females). All participants provided informed consent and the data collection was approved by the Psychology Research Ethics Committee at the Institute of Psychology at Leiden University.

### Smartphone behavioral recording and the joint-interval distribution

A background app (TapCounter, QuantActions AG) was used to record the time-stamp of all smartphone touchscreen interaction events^41^. The timestamp of the event onset – say towards a swipe or a tap – was recorded in UTC milliseconds. The data was collected using a unique participant identifier linking to the self-reported age, and gender. The background app also logged the label of the App in use but this information was not used here. Apart from the touchscreen events, the App also recorded the screen “on” and “off” events which were used to define within-session interactions.

Within the session (between a “screen on” event and a “screen off” event) inter-touch intervals (ITIs) were used to estimate joint-interval distributions. The intervals were accumulated in consecutive 1-hour bins. For each hourly bin, the duration of an interval with index *k* was related to that of the subsequent interval *k* + 1, thus creating a two-dimensional map of events. We consider each subsequent pair of events as being sampled from a joint-probability distribution *P*(*k, k* + 1), and this estimated the continuous two-dimensional joint probability distribution over the log_10_ transformed space using kernel density estimation with a Gaussian kernel and a bandwidth of 0.1 (KernelDensity, sklearn, python). We limited our range of estimation between 10^1.5^ ms (∼30ms) to 10^5^ ms (∼2 min), with the upper bound corresponding to the 99^th^ percentile of all observed inter-events. We discretize the space in 50 steps in each dimension thus obtaining a behavioral space of 50 × 50 x *D*, where *D* is the recording duration in hours.

### Wavelet transformation and block-bootstrap

The continuous wavelet transform (*cwt*, MATLAB, MathWorks, Natick) of the time-series of each two-dimensional bin of the JID was based on a filter bank (*cwtfilterbank*, MATLAB) with the following parameters: sampling period of 1h, voices per octave of 40, and period limits from 2 hours to 80 days. The cone of influence was removed before estimating the periodogram power (*P*) at a given period (*t*), where the matrix *S*_*i*_ contained the wavelet transformed spectral values:

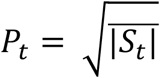

The time series was block-bootstrapped with a 24-hour window, and the wavelet transform was estimated for each bootstrap (number of targeted bootstraps 1000)^27^. The bootstrap computations were performed on a high-performance computing cluster (ALICE, Leiden University). The computations failed in 11 of the participants, resulting in an N of 401. The mean of the bootstrapped values was used to capture the aperiodic component of the periodogram, and it was subtracted from the real to obtain the adjusted periodogram (power index).

The significant periodogram values were isolated based on a two-tailed *α* = 0.05 based on the bootstrapped periodogram. This was subsequently corrected for multiple comparisons using clustering (LIMO EEG^30^) across the JID feature space and time such that a cluster contained significant values in 5 neighboring bins. Briefly, the cluster sizes were accumulated for each boot against the rest of the boots, and the maximum cluster size was noted for each iteration. Only those clusters based on real data which were > 97.5^th^ percentile of the iterated set of maximum clusters were considered significant.

### Non-negative matrix factorization

For each individual the spectral computations using wavelet transform resulted in a tensor of the dimension of the JID (50 × 50) x *T*, where *T* is the number of periods in the periodogram and is a function of the recording length *R* (in the population min. *T* was 335, max. was 396). The bootstrap-derived aperiodic component was subtracted from this tensor. The tensor was then sliced along the periods dimension to select only the periods between *T*_*min*_(with index 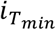 in the periods list) and *T*_*max*_(with index 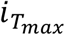 in the period list) reshaped into a matrix of sizes *T*′ × 2500 where 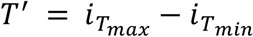. The matrix was then globally shifted up by subtracting the global minimum rendering it non-negative.

The non-negative matrix factorization (*mexTrainDL, spams package for MATLAB, Inria*) yielded two matrices: a meta-rhythms matrix *W* ∈ ℝ^*T*′×*R*^ and a meta-behavior matrix *H* ∈ ℝ^*R*×2500^ where R is the optimal factorization rank. The optimal rank for the factorization was estimated using cross-validation. For the cross-validation, we repeated the factorization for ranks from 3 to 15. We randomly selected 10% of the entries from the matrix to be masked and used as a test set. The factorization was done on the matrix with masked values. After the factorization, the test error was obtained by evaluating the reconstruction error of the 10% of entries that were left out (test set). We repeated this process 100 times for each rank. The optimal rank was defined as the rank with the minimum mean test error across the 100 repetitions. The optimal rank varied from person to person with a minimum rank of 3 and a maximum of rank 14. The factorization was performed for all the subjects (N = 401) with *P*_*min*_= 2.2 days and *P*_*max*_= 27.7 days 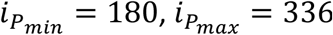, thus *T*^′^= 147. And separate factorization was performed for subjects with at least 180 days of data (N = 218) with *P*_*min*_= 2.2 days and *P*_*max*_= 70.5 days, 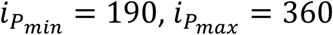, thus *T*^′^= 201. Note, although the seven individuals who did not report their gender were included in the factorization but they were omitted from the plots.

While non-negative matrix factorization, is a powerful dimensionality reduction method, it is known to be susceptible to its initial conditions and can lead to different factorizations if repeated with the same input matrix ^42^. To mitigate this issue, for each individual we repeated the following procedure to pick the most reproducible (as defined below) decomposition: Given the best rank calculated with the procedure described above, we repeated the decomposition 1000 times with different initial conditions. We then calculated the pairwise correlation coefficient for each pair of decompositions (using the meta-rhythms matrix) by calculating the cross-correlation matrix across components and picking the highest correlation across the components (meta-rhythms). For each decomposition, we then computed the median of the pairwise cross-correlation with all other decompositions and picked as the final decomposition the one with the highest median correlation across the 1000 repetitions.

We clustered the meta-rhythms accumulated across the population by using a one-dimensional continuous wavelet transform (*mdwtcluster*, MATLAB). The one-dimensional CWT clustering uses all the coefficients from a Daubechies 4 wavelet. We first *z*-scored each meta-rhythm. Then we estimated the optimal number of clusters by using the silhouette method (*evalclusters*, MATLAB). Finally, we operationalized the clustering with the obtained best number of clusters. The best number of clusters emerged to be 8 for the analysis up to 27.7 days and 9 for the analysis up to 70.5 days. Note, that since we considered each meta-rhythm independently and each subject had a different number of meta-rhythms, each individual could contribute with zero to multiple meta-rhythms to a given cluster.

### Phase coherence analysis

We selected 139 subjects with at least 180 days of overlapping recordings. We then extracted a range of periods for each cluster corresponding to the periods surrounding the peak of the mean meta-rhythm across the cluster. More specifically, for each cluster, we first calculated the mean across the cluster. The location of the peak of the mean was thus identified as the reference period for that meta-rhythm cluster. To get the ranges for each peak (min/max range), we first estimated the 95^th^ percentile of the mean of the cluster, and then we created a binary vector indicating where the values of the mean cluster were > 95^th^ percentile, the min. the range is the period corresponding to the minimum index that has a 1 in the above binary vector, while the max. the range is the period corresponding to the maximum index that has a 1 in the above binary vector.

For each subject and each period range, we selected the JID two-dimensional bin with the maximum power index located in the largest statistical cluster (largest in terms of the overall number of significant pixels for a select period). For each pair of subjects (thus selected JID) we extracted the probability density of that pixel from hourly JIDs. We calculated the CWT spectral coherence based on the same filterbank and wavelet described above (*wcoherence*, MATLAB). Over the spectral coherence spectrum, we selected the period range of interest and estimated the average coherence. This was repeated for all possible pairs and at every discovered multi-day rhythm. The coherence values above 95^th^ percentile (corresponding to one-tail α = 0.05) of 24 h block bootstrapped data (∼ 1000 bootstraps as mentioned above) were considered significant.

## Acknowledgment

This study was funded by Velux Stiftung (project no. 1283, awarded to AG). EC was supported by the SNSF Early Postdoc.Mobility (no. 199692 awarded to EC with AG as host). The authors would like to thank the student and staff researchers behind the data collection platform agestudy.nl. In particular, we would like to thank Guido Band for his constructive edits on this manuscript.

## Code availability

The codes written specifically for this analysis are shared here: https://github.com/codelableidenvelux/Multi_dayOscillations_2022

## Data availability

The periodograms and the corresponding bootstrapped statistics are shared on dataverse.nl (temporary data link for review: https://surfdrive.surf.nl/files/index.php/s/GeESDy5lfY4q17j. The bootstrapped periodograms are also available upon request (but not shared due to the high data volume).

## Related Supplementary Figures

Supplementary Figure 1: C*lusters of meta-rhythms across the population, obtained with individualized optimal-rank non-negative matrix factorization for individuals based on the data collections with at least 180 days of recordings*. The panels are the same as in Figure 3, but here all the clusters obtained from the analysis with Periods up to 70 days are shown.

Supplementary Figure 2: C*lusters of common meta-rhythms across the population, obtained with individualized optimal-rank non-negative matrix factorization for the individuals based on the data collections with at least 90 days of recordings*. The panels are the same as in Figure 3, but here all the clusters obtained from the analysis with Periods up to 28 days are shown.

Supplementary Figure 3: Meta-behaviors according to the factorization involving at least 180 days of recording. Top row: The mean meta-behaviors are shown for each of the identified multi-day rhythms (including all of the subjects with a given meta-rhythm). Middle row: The mean-meta behaviors based on the individuals where the identified rhythm was part of a statistically significant cluster according to the parametric statics based on block bootstraps. Bottom row: The probability of observing a statistically significant periodogram deflection according to parametric statistics in the period ranges indicated in ‘[]’.

Supplementary Figure 4: Meta-behaviors according to the factorization involving at least 90 days of recording. Top row: The mean meta-behaviors are shown for each of the identified multi-day rhythms (including all of the subjects with a given meta-rhythm). Middle row: The mean-meta behaviors based on the individuals where the identified rhythm was part of a statistically significant cluster according to the parametric statics based on block bootstraps. Bottom row: The probability of observing a statistically significant periodogram deflection according to parametric statistics in the period ranges indicated in ‘[]’.

Supplementary Figure 5: *Widespread coherence across the population for diurnal (24h) rhythm*. The spectral coherence for the 24-hours rhythm across individuals.

## References

1. Dijk, D.-J. & Archer, S. N. Light, Sleep, and Circadian Rhythms: Together Again. PLOS Biol. 7, e1000145 (2009).

2. Wager-Smith, K. & Kay, S. A. Circadian rhythm genetics: from flies to mice to humans. Nat. Genet. 26, 23–27 (2000).

3. Mistlberger, R. E. & Skene, D. J. Social influences on mammalian circadian rhythms: animal and human studies. Biol. Rev. 79, 533–556 (2004).

4. Schmidt, C., Collette, F., Cajochen, C. & Peigneux, P. A time to think: Circadian rhythms in human cognition. Cogn. Neuropsychol. 24, 755–789 (2007).

5. Casiraghi, L. et al. Moonstruck sleep: Synchronization of human sleep with the moon cycle under field conditions. Sci. Adv. 7, eabe0465 (2021).

6. Cordi, M. et al. Lunar cycle effects on sleep and the file drawer problem. Curr. Biol. 24, R549–R550 (2014).

7. Cajochen, C. et al. Evidence that the Lunar Cycle Influences Human Sleep. Curr. Biol. 23, 1485–1488 (2013).

8. James, A. The validity of ‘;biorhythmic’ theory questioned. Br. J. Psychol. 75, 197–200 (1984).

9. Persinger, M. A., Cooke, W. J. & Janes, J. T. No Evidence for Relationship between Biorhythms and Industrial Accidents. Percept. Mot. Skills 46, 423–426 (1978).

10. Peveto, N. The Relationship of Biorhythms to Academic Performance in Reading. LSU Hist. Diss. Theses (1980) doi:10.31390/gradschool_disstheses.3577.

11. Owen, C., Tarantello, C., Jones, M. & Tennant, C. Lunar Cycles and Violent Behaviour. Aust. N. Z. J. Psychiatry 32, 496–499 (1998).

12. Laverty, W. H. & Kelly, I. W. Cyclical Calendar and Lunar Patterns in Automobile Property Accidents and Injury Accidents. Percept. Mot. Skills 86, 299–302 (1998).

13. Ichino, A. & Moretti, E. Biological Gender Differences, Absenteeism, and the Earnings Gap. Am. Econ. J. Appl. Econ. 1, 183–218 (2009).

14. Shansky, R. M. Are hormones a “female problem” for animal research? Science 364, 825–826 (2019).

15. Pletzer, B., Harris, T.-A., Scheuringer, A. & Hidalgo-Lopez, E. The cycling brain: menstrual cycle related fluctuations in hippocampal and fronto-striatal activation and connectivity during cognitive tasks. Neuropsychopharmacology 44, 1867–1875 (2019).

16. Clare, A. W. Invited review hormones, behaviour and the menstrual cycle. J. Psychosom. Res. 29, 225–233 (1985).

17. Baud, M. O. et al. Multi-day rhythms modulate seizure risk in epilepsy. Nat. Commun. 9, 88 (2018).

18. Karoly, P. J. et al. Cycles in epilepsy. Nat. Rev. Neurol. 1–18 (2021) doi:10.1038/s41582-021-00464-1.

19. Leguia, M. G. et al. Seizure Cycles in Focal Epilepsy. JAMA Neurol. 78, 454–463 (2021).

20. Wehr, T. A. Bipolar mood cycles and lunar tidal cycles. Mol. Psychiatry 23, 923–931 (2018).

21. Wehr, T. A. & Helfrich-Förster, C. Longitudinal observations call into question the scientific consensus that humans are unaffected by lunar cycles. BioEssays 43, 2100054 (2021).

22. Karoly, P. J. et al. Multiday cycles of heart rate are associated with seizure likelihood: An observational cohort study. eBioMedicine 72, (2021).

23. Huber, R. & Ghosh, A. Large cognitive fluctuations surrounding sleep in daily living. iScience 24, 102159 (2021).

24. Golder, S. A. & Macy, M. W. Diurnal and Seasonal Mood Vary with Work, Sleep, and Daylength Across Diverse Cultures. Science 333, 1878–1881 (2011).

25. Leise, T. L. Wavelet analysis of circadian and ultradian behavioral rhythms. J. Circadian Rhythms 11, 5 (2013).

26. Ceolini, E. et al. A model of healthy aging based on smartphone interactions reveals advanced behavioral age in neurological disease. iScience 25, 104792 (2022).

27. Cazelles, B., Cazelles, K. & Chavez, M. Wavelet analysis in ecology and epidemiology: impact of statistical tests. J. R. Soc. Interface 11, 20130585 (2014).

28. Lee, D. D. & Seung, H. S. Learning the parts of objects by non-negative matrix factorization. Nature 401, 788–791 (1999).

29. Brunet, J.-P., Tamayo, P., Golub, T. R. & Mesirov, J. P. Metagenes and molecular pattern discovery using matrix factorization. Proc. Natl. Acad. Sci. 101, 4164–4169 (2004).

30. Pernet, C. R., Chauveau, N., Gaspar, C. & Rousselet, G. A. LIMO EEG: A Toolbox for Hierarchical LInear MOdeling of ElectroEncephaloGraphic Data. Computational Intelligence and Neuroscience https://www.hindawi.com/journals/cin/2011/831409/ (2011) doi:10.1155/2011/831409.

31. Karoly, P. J. et al. Epileptic Seizure Cycles: Six Common Clinical Misconceptions. Front. Neurol. 12, (2021).

32. Bachleitner, W., Kempinger, L., Wülbeck, C., Rieger, D. & Helfrich-Förster, C. Moonlight shifts the endogenous clock of Drosophila melanogaster. Proc. Natl. Acad. Sci. 104, 3538–3543 (2007).

33. Bernard, C. Circadian/multidien Molecular Oscillations and Rhythmicity of Epilepsy (MORE). Epilepsia 62, S49–S68 (2021).

34. Novák, B. & Tyson, J. J. Design principles of biochemical oscillators. Nat. Rev. Mol. Cell Biol. 9, 981–991 (2008).

35. Lugo, C. A. & McKane, A. J. Quasicycles in a spatial predator-prey model. Phys. Rev. E 78, 051911 (2008).

36. Nisbet, R. M. & Gurney, W. S. C. A simple mechanism for population cycles. Nature 263, 319–320 (1976).

37. Esmaeili, S., Hastings, A., Abbott, K. C., Machta, J. & Nareddy, V. R. Noise-induced versus intrinsic oscillation in ecological systems. Ecol. Lett. 25, 814–827 (2022).

38. Barraquand, F. et al. Moving forward in circles: challenges and opportunities in modelling population cycles. Ecol. Lett. 20, 1074–1092 (2017).

39. Zwan, M. D. et al. Dutch Brain Research Registry for study participant recruitment: Design and first results. Alzheimers Dement. Transl. Res. Clin. Interv. 7, e12132 (2021).

40. Borger, J. N., Huber, R. & Ghosh, A. Capturing sleep–wake cycles by using day-to-day smartphone touchscreen interactions. Npj Digit. Med. 2, 1–8 (2019).

41. Balerna, M. & Ghosh, A. The details of past actions on a smartphone touchscreen are reflected by intrinsic sensorimotor dynamics. Npj Digit. Med. 1, 4 (2018).

42. Wu, S. et al. Stability-driven nonnegative matrix factorization to interpret spatial gene expression and build local gene networks. Proc. Natl. Acad. Sci. 113, 4290–4295 (2016).

